# Variability in drought gene expression datasets highlight the need for community standardization

**DOI:** 10.1101/2024.02.04.578814

**Authors:** Robert VanBuren, Annie Nguyen, Rose A. Marks, Catherine Mercado, Anna Pardo, Jeremy Pardo, Jenny Schuster, Brian St. Aubin, Mckena Lipham Wilson, Seung Y. Rhee

## Abstract

Physiologically relevant drought stress is difficult to apply consistently, and the heterogeneity in experimental design, growth conditions, and sampling schemes make it challenging to compare water deficit studies in plants. Here, we re-analyzed hundreds of drought gene expression experiments across diverse model and crop species and quantified the variability across studies. We found that drought studies are surprisingly uncomparable, even when accounting for differences in genotype, environment, drought severity, and method of drying. Many studies, including most Arabidopsis work, lack high-quality phenotypic and physiological datasets to accompany gene expression, making it impossible to assess the severity or in some cases the occurrence of water deficit stress events. From these datasets, we developed supervised learning classifiers that can accurately predict if RNA-seq samples have experienced a physiologically relevant drought stress, and suggest this can be used as a quality control for future studies. Together, our analyses highlight the need for more community standardization, and the importance of paired physiology data to quantify stress severity for reproducibility and future data analyses.

## Introduction

Drought, increasingly prevalent in both natural and agricultural landscapes, is escalating in frequency and severity due to the dynamic climate. This trend has spurred the development of an extensive and increasingly interdisciplinary research community focused on understanding plant adaptation to water-limited environments (Osmolovskaya et al., 2018; Ekundayo et al., 2022). Meteorologically, drought manifests as drier than normal conditions, but its physiological impact on plants varies based on the duration, severity, and timing of the stress events, alongside local soil and habitat conditions (Gupta et al., 2020; Tardieu, 2012). Mild, infrequent drought events may result in only slight reductions in photosynthesis and growth, often without significant impacts on biomass or yield. In contrast, recurrent or severe bouts of drought may cause unrecoverable damage or even plant death (Farooq et al., 2009). Central to understanding and engineering drought resilience is the ability to apply consistent, physiologically relevant, and reproducible stress events across scales (Großkinsky et al., 2015). Such standardization is necessary to develop a community framework that allows for comparison and expansion of previous experiments (Juenger and Verslues, 2022).

Water deficit responses likely evolved during terrestrialization, and they have been continually refined, repurposed, and diversified to enable plants to colonize virtually every biome (Bowles et al., 2021). Resistance to drought is an emergent phenotype involving the synchronization of numerous physiological and genetic processes, and diverse lineages of plants have evolved numerous adaptations to avoid, escape, and tolerate water deficits (Artur and Kajala, 2021);(Chaves and Oliveira, 2004; Turner, 1986). Different plant lineages, populations, or even individual genotypes use combinations of these strategies to tolerate water limitations (Verslues and Juenger, 2011; Basu et al., 2016; Farooq et al., 2009). The genetic mechanisms underlying responses or tolerance to drought stress are highly complex and involve the activation of hundreds to thousands of pathways that collectively enable resilience to water deficit. Most drought related pathways were discovered and characterized in the model plant Arabidopsis, but core regulatory, biochemical, and physiological responses are broadly conserved across green plants (Shinozaki and Yamaguchi-Shinozaki, 2007). The genetic basis of adaptations to water deficit is an active and exciting area of plant science research, and numerous important research gaps still remain (Verslues et al., 2023; Eckardt et al., 2023).

One promising approach to closing the knowledge gaps in understanding the genetic basis of drought adaptation is using large-scale omics technologies. Numerous large-scale datasets have been collected across diverse plant lineages to study the effects of drought stress. Some studies have measured physiological responses in naturally water limited environments (Pardo et al., 2022; Danilevskaya et al., 2019; Groen et al., 2022), but most use simulated drought events under controlled or semi-controlled conditions to induce water deficit responses (Gonzalez et al., 2022). Simulated drought studies range in scale and severity from large rainout shelters withholding water from thousands of plants in an ecological or agricultural setting to agar plates containing solutes to lower water potential. Each of these approaches has benefits and drawbacks related to cost, consistency, and accuracy of applying drought. For example, using polyethylene glycol, mannitol, and salt to lower water potential may not actually induce true drought responsive pathways (Gonzalez et al., 2022), and restricting plants to small pots in growth chambers or greenhouses can impact root growth and lead to physiologically irrelevant and irreproducible drying (Granier et al., 2006). Individual labs utilize radically different experimental approaches, growth conditions, and sampling schemes for drought assays, and these added variables mask emergent properties of an already complex phenotype. A major challenge for cross-species analysis is finding comparable biological datasets with similar design, implementation, and sampling.

Here, to evaluate the comparability and reusability of drought gene expression data, we compared public datasets across labs and experiments, and searched for patterns that delineate drought and control conditions. We first focused on data from the model plant *Arabidopsis thaliana* and then expanded our analyses to include five additional model and crop species with the most published drought data. We found that drought gene expression data are more variable compared to other abiotic stresses, and many studies lack basic physiological data to assess the magnitude or even presence of water deficit stress. Our analyses highlight the need for community standardization to enable the reuse and integration of datasets across scales from different laboratories.

## Results

### Variability of gene expression datasets on drought in Arabidopsis

Plant responses to drought is a well-studied topic. There are ∼34,000 articles in PubMed related to drought stress in plants, and this wealth of knowledge has uncovered numerous phenotypes, pathways, and genes underlying responses to water deficit (Gupta et al., 2020). Most of our understanding of drought responses at the molecular genetic level is based on work in Arabidopsis, including over 100 studies surveying genome-wide gene expression (RNAseq) changes under water deficit across different accessions, environmental conditions, and mutant backgrounds (Supplemental Table 1). Collectively, these datasets have been incorporated into public gene expression atlases, co-expression networks, and other tools that are broadly used by the plant science community to understand which genes and pathways underlie drought responses (Lamesch et al., 2012; Klepikova et al., 2016). However, drought experiments vary wildly in the degree, severity, and implementation of water deficit. Many experiments are analyzed in isolation and arrive at independent conclusions. The choice of which drought experiments to reference for future studies can drastically alter hypothesis generation and inference of biological function. This raises a fundamental question, how comparable are drought studies across different experiments?

To survey the variability in public drought data, we re-analyzed 109 water deficit RNAseq experiments in Arabidopsis obtained from the Sequence Read Archive (SRA). We manually curated metadata from 1,301 RNAseq samples across these 109 BioProjects. These datasets include a range of genotypes and mutant backgrounds, developmental time points and tissues, and differences in stress severity and duration across a range of natural or controlled drought conditions (Figure 1; Supplemental table 1). Based on the available metadata, 81% of studies were conducted in growth chambers in standard potting media, 13% on agar plates, and 5% in greenhouses. Half of the studies (51%) applied a natural dry down by stopping irrigation, 27% had controlled drying to a set soil moisture content, 8% removed plants from media and let them air dry, and 14% used PEG to lower water potential or ABA to simulate water deficit responses. Surprisingly, 39% of Arabidopsis studies did not report paired physiology data measuring plant stress, such as gas exchange, photosynthesis, leaf water potential, or leaf relative water content. Next, we processed the raw reads through a common pipeline to remove variation arising from the different algorithmic and statistical frameworks used in each individual study.

**Figure 1.**
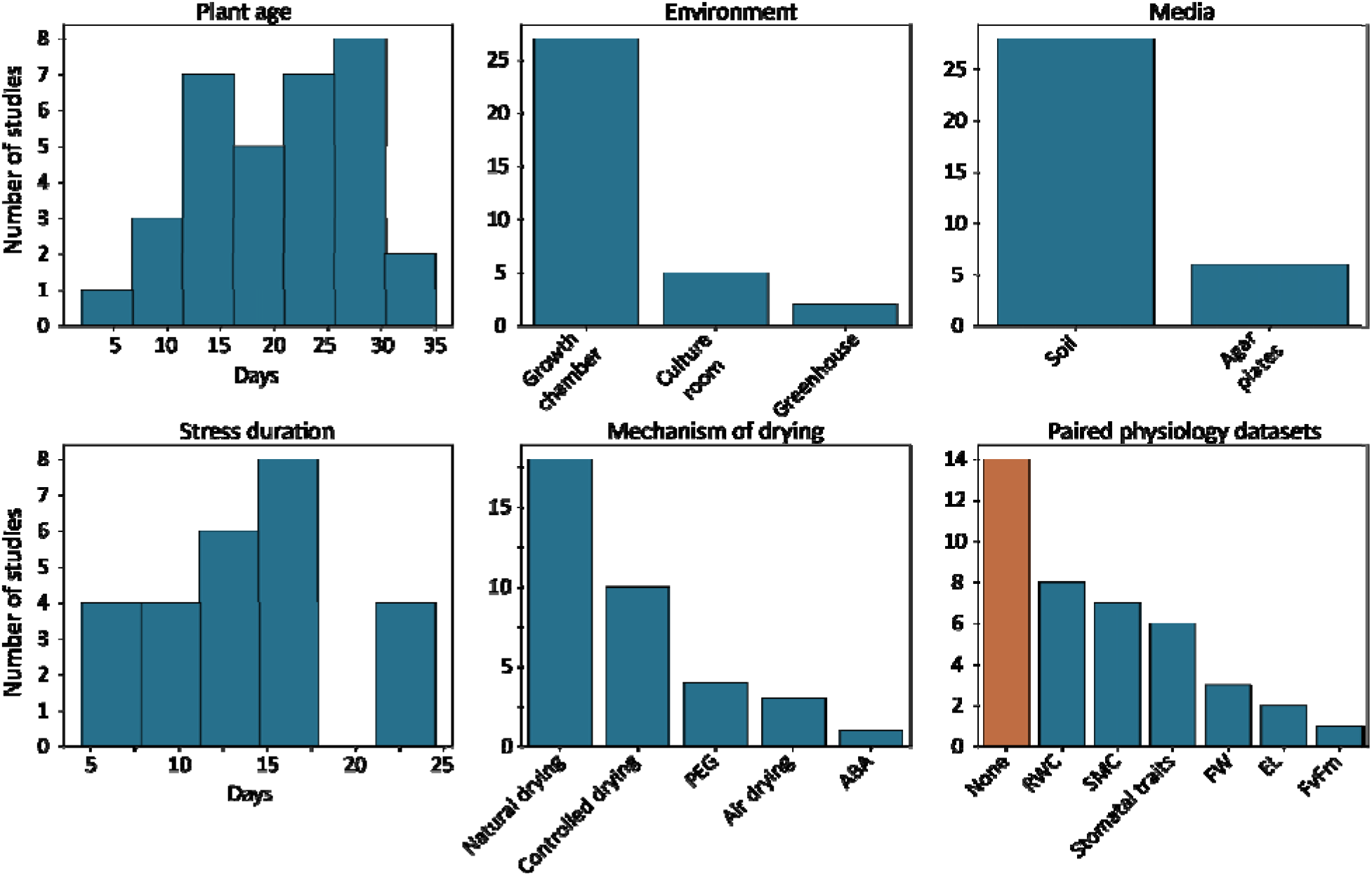
Summary of Arabidopsis drought gene expression metadata. Metadata was collected for the 36 BioProjects with associated publications. The top row shows a histogram of developmental stage or age of plants (in days) at sampling, the environment where studies were conducted, and the media plants were propagated in. The bottom row shows a histogram of the duration of water deficit stress, mechanism of drying, and paired physiology data. Studies using PEG, air drying, and ABA are not plotted in the stress duration graph, as experiment times ranged from 1-8 hours. Abbreviations are as follows: relative water content (RWC), soil moisture content (SMC), fresh weight (FW), and electrolyte leakage (EL).

Raw Illumina RNAseq reads were quality trimmed and aligned to the TAIR10 gene models, and raw or batch corrected expression values in transcripts per million (TPM) were used as a basis for downstream analysis.

To identify any factors that clearly delineate samples within or across experiments, we used dimensionality reduction with an expectation that samples should cluster by water stress status. Principal component analysis (PCA) and t-Distributed Stochastic Neighbor Embedding (t-SNE) show no clear separation between drought-treated and control samples across Arabidopsis drought experiments (Figure 2a; Supplemental Figure 1, 2). Within experiments, some BioProjects show clear separation of drought and control samples/replicates, but a surprising number have interspersed samples (Supplemental Figure 3). Across experiments, samples were broadly separated by tissue type and BioProject. Root, seedlings, inflorescence, and siliques form groups in a similar dimensional space, whereas clusters of leaf and whole plant samples were more dispersed across PC1 and PC2 or dimensions 1 and 2 for PCA and t-SNE, respectively (Figure 2a, Supplemental Figure 2). Samples within the same experiment or BioProject tended to cluster together, suggesting that the experimental effects are responsible for much of the separation we observed (Supplemental Figure 3). Individual accessions of Arabidopsis show remarkable differences in expression dynamics under drought (Des Marais et al., 2012), and we tested if genotype differences may explain the lack of correlation between stressed samples across experiments. Similar to other experimental factors, there is no clear separation by accession (Figure 2a). There is also no clear separation of samples by the reported duration, severity, or type of water deficit (e.g., drying vs solute based), or technical variables such as sequencing read length, technology, read chemistry, or publication year.

**Figure 2.**
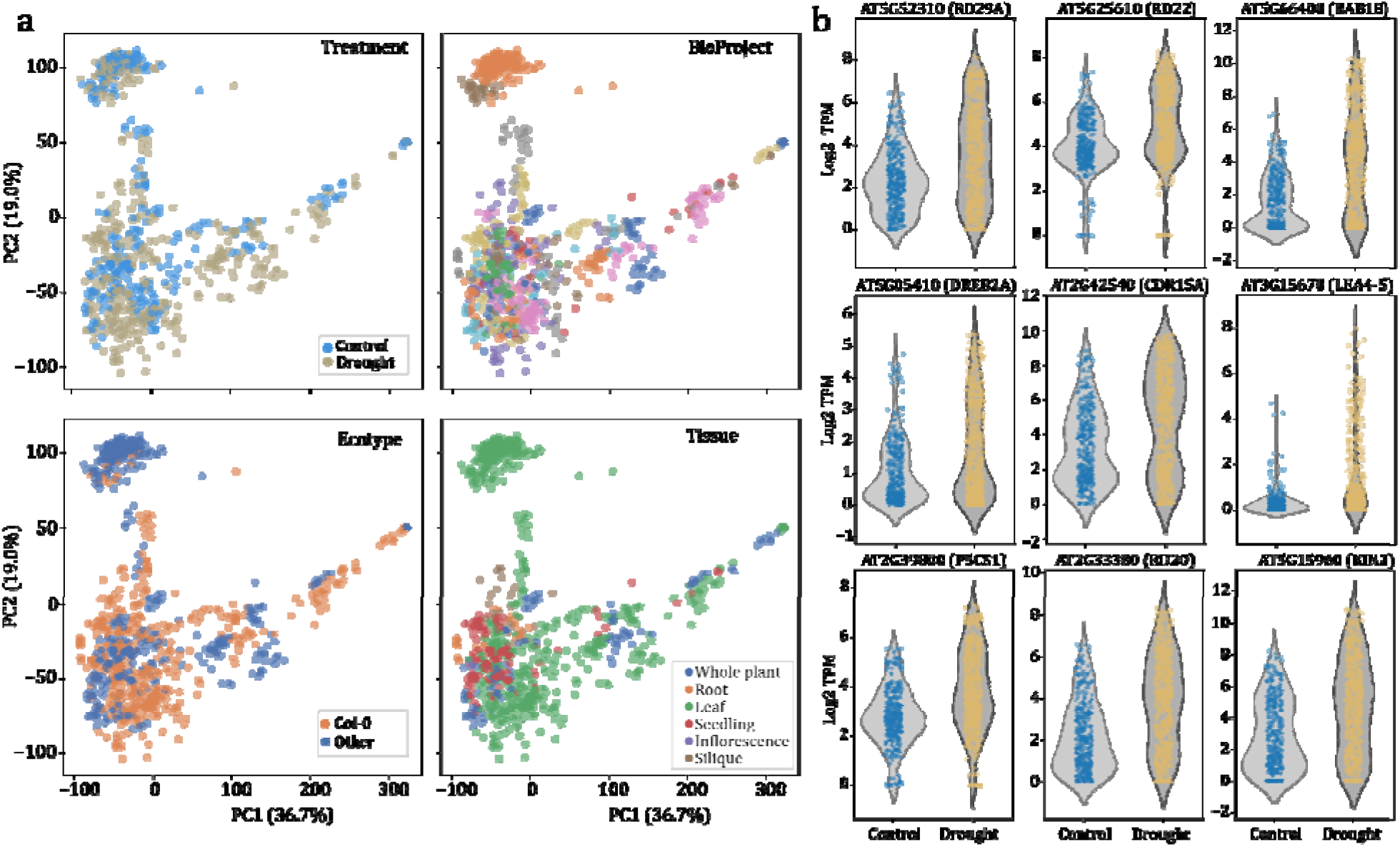
Dramatic variability of public drought gene expression datasets in Arabidopsis. (a) Principal component analysis of 1,301 drought related RNAseq samples collected from the sequence read archive (SRA). The first two principal components are plotted for all samples and colored by different factors including a binary classification of drought and control (upper left), BioProject (upper right), genotype/accession of the sample (Col-0 or others; bottom left), and the tissue type (bottom right). (b) Violin plot of log2 transformed TPM of RNAseq data for nine drought marker genes in samples classified as control (left) and drought (right).

To test if the observed separation by experiment (BioProject) rather than stress-related factors is caused by batch effects, we applied ComBat (Behdenna et al., 2023) to the expression matrix. This method leverages an empirical Bayes framework to estimate and subsequently adjust for batch effects. Post-ComBat adjustment, the first two principal components of the expression values accounted for only 19% of the total variance (Supplemental Figure 4). Even when batch effects associated with BioProject were addressed, the samples did not show clear differentiation between stress and non-stress conditions, and much of the variance that was removed relates to true biological differences rather than technical artifacts. Together, this suggests drought experiments in Arabidopsis have extreme variability and individual datasets are largely incomparable using traditional approaches.

We next sought to understand why the Arabidopsis drought RNAseq data appeared so variable across experiments. We first hypothesized that since dimensionality reduction provides a summary across all genes, individual factors such as drought stress may be confounded by other experimental factors. Therefore, to see if confounding factors are masking clustering of drought-stressed samples, we surveyed the expression pattern of nine drought marker genes across all the samples. If confounding factors indeed masked the drought-response programs of relevant genes, we would expect the drought-marker genes to be consistently induced in drought across all experiments. The nine drought marker genes included RD29A (Nordin et al., 1991), RD22 (Yamaguchi-Shinozaki and Shinozaki, 1993), RAB18, DREB2A (Liu et al., 1998), COR15A (Hincha et al., 2021), LEA4-5 (Bray, 2004), P5CS1 (Yoshiba et al., 1995), RD20, and KIN2. Marker genes were generally expressed at higher levels in drought-stressed samples compared to well-watered, but this is highly variable across our datasets (Figure 2b, Supplemental Figure 5). For instance, the dehydrin RAB18 and ABA induced transcription factor DREB2 were highly expressed under drought but generally not expressed under well-watered conditions. However, roughly a third of samples labeled as ‘drought’ had no detectable expression of these two genes (Figure 2b, Supplemental Figure 5). This pattern is consistent across all nine drought marker genes, suggesting the hypothesis that confounding factors are masking the drought response program is not supported. Instead, the data show that many drought-treated samples lack a molecular signature of water deficit, and may not have experienced a physiologically relevant drought stress.

We then wondered whether the observed inconsistencies in drought expression data might simply reflect a wide range of responses to various drought regimes applied, given the varying drought intensities and diverse growing conditions of different experiments. To understand if a similar variability exists in response to other stresses, we re-analyzed publicly available expression datasets related to heat stress in Arabidopsis. There are fewer published heat expression datasets, and we curated 21 experiments where plants were subjected to physiologically relevant heat stress in Arabidopsis between temperature ranges of 35-42 C. Metadata indicated that growth conditions were similarly variable as drought studies, and the raw reads were processed as described above. Strikingly, when we performed dimensionality reduction on log-transformed expression data, the samples were distinctly categorized into either ‘heat’ or ‘control’ groups based on principal component 2, which explained 14% of the variation (Figure 3a). Similar to the drought dataset, principal component 1, which explained 21% of the variation, differentiated the samples based on tissue type, grouping them as whole seedlings or leaves. Unlike in the drought dataset, there was no distinct separation by experiment (BioProject). We surveyed the expression patterns of four heat marker genes (HSP70, HSP90, MBF1c, and DREB2A) to see if individual genes have a similarly clear pattern. Heat marker genes have notably higher expression in all heat stressed samples compared to control, and there is little overlap in the distribution of expression levels of these marker genes between the two conditions (Figure 3b, 3c). Thus, public heat stress expression datasets demonstrate a clear molecular signature of heat stress. This signature persists through any experimental variation across studies, indicating that the inconsistencies noted in the drought stress data are not simply artifacts of work conducted in different laboratories.

**Figure 3.**
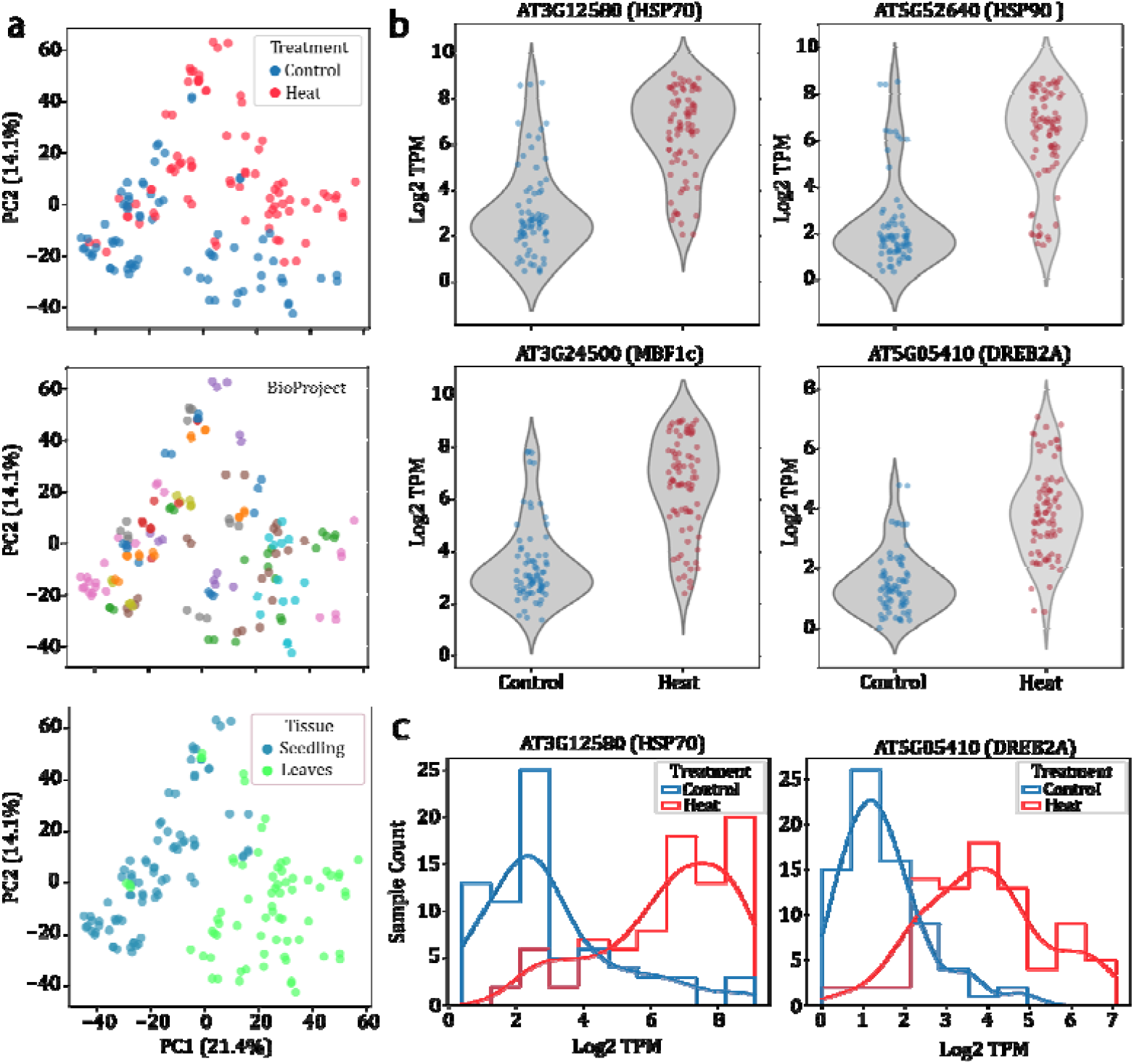
Comparison of public heat stress gene expression data in Arabidopsis. (a) Principal component analysis of 156 heat stress related RNAseq samples collected from the sequence read archive (SRA). The first two principal components are plotted for all samples and colored by different factors including a binary classification of heat stressed and control (top), BioProject (middle), and the tissue type (bottom). (b) Violin plot of log2 transformed TPM of RNAseq data for four drought marker genes in samples classified as control (left) and heat stress (right). (c) Histogram of the same log2 transformed gene expression data as (b) for two heat stress marker genes of HSP70 (At3G12580; left), and DREB2A (AT5G05410; right).

### Developing a predictive model for classifying drought gene expression

Dimensionality reduction and clustering approaches were unable to delineate drought-stressed from control samples, and we hypothesize this was driven by quality issues of the underlying datasets. We previously developed a cross-species predictive model to accurately classify drought stressed RNAseq data in maize and sorghum (Pardo et al., 2022), and sought to test if this approach could differentiate among drought and control samples in Arabidopsis. We developed Random Forest (RF) based predictive models to classify the Arabidopsis samples as “drought” or “control” based on normalized gene expression values alone. We divided the RNAseq samples into a training set with 75% of the experiments (BioProjects) and a testing set with the remaining 25%. The overall accuracy of our predictive model was 66% (Supplemental Table 2), which is substantially lower than the model developed with high-quality maize and sorghum data (Pardo et al., 2022). The precision and recall for ‘drought’ samples were 0.79 and 0.51, respectively and 0.61 and 0.84 for control. We tested four other classifiers including Linear Support Vector Classifier (SVC), Simple Neural Network (MLP), Histogram-based Gradient Boosting Classifier (HGB), and K-Nearest Neighbor Classifier (KNN) to see if this improved predictive accuracy. HGB had similar performance to RF (overall accuracy 65%) but SVC, KNN, and MLP performed significantly worse with 52%, 52%, and 56% overall accuracy (Supplemental Table 2). The relatively low predictive accuracy in Arabidopsis was initially surprising, as our model was trained with significantly more data and tested within a single species with less genetic diversity than either maize or sorghum. Further, Arabidopsis experiments are generally conducted within a narrower set of conditions compared to the diverse growth chamber, greenhouse and field environments for maize and sorghum. We suspect that the reduced predictive accuracy in Arabidopsis might be due to the inclusion of datasets with plants that are not physiologically stressed, thereby diminishing the effectiveness of our models.

To assess the efficiency of the RF models in predicting drought and control samples for each Arabidopsis experiment, we employed the Leave-One-Group-Out cross-validation method. This method involved iterative training of the model using all datasets except one, and then gauging both its overall and individual sample performances. The aggregate performance was 0.71, with a precision of 0.77 and a recall of 0.75. Performance metrics for individual datasets varied widely, from a purely random prediction (approximately 0.5) to absolute accuracy (1.0) (Figure 4d). Random Forest had perfect or near perfect prediction in six experiments, but notably, 14 Arabidopsis experiments had performances nearing random results, with scores less than 0.65. Three of these experiments reported ‘mild’ drought stress events where the authors collected limited physiology data to support the degree of plant stress (Dubois et al., 2017)(Clauw et al., 2016, 2015). This includes two large-scale experiments of 6 and 98 Arabidopsis accessions subjected to ‘mild’ drought stress in an automated plant phenotyping platform (Clauw et al., 2016, 2015). These experiments collected limited physiology data aside from growth rate and soil moisture content, making it difficult to evaluate if the plants were experiencing a physiologically relevant drought stress event, or if they were simply growing with less but sufficient soil water content.

**Figure 4.**
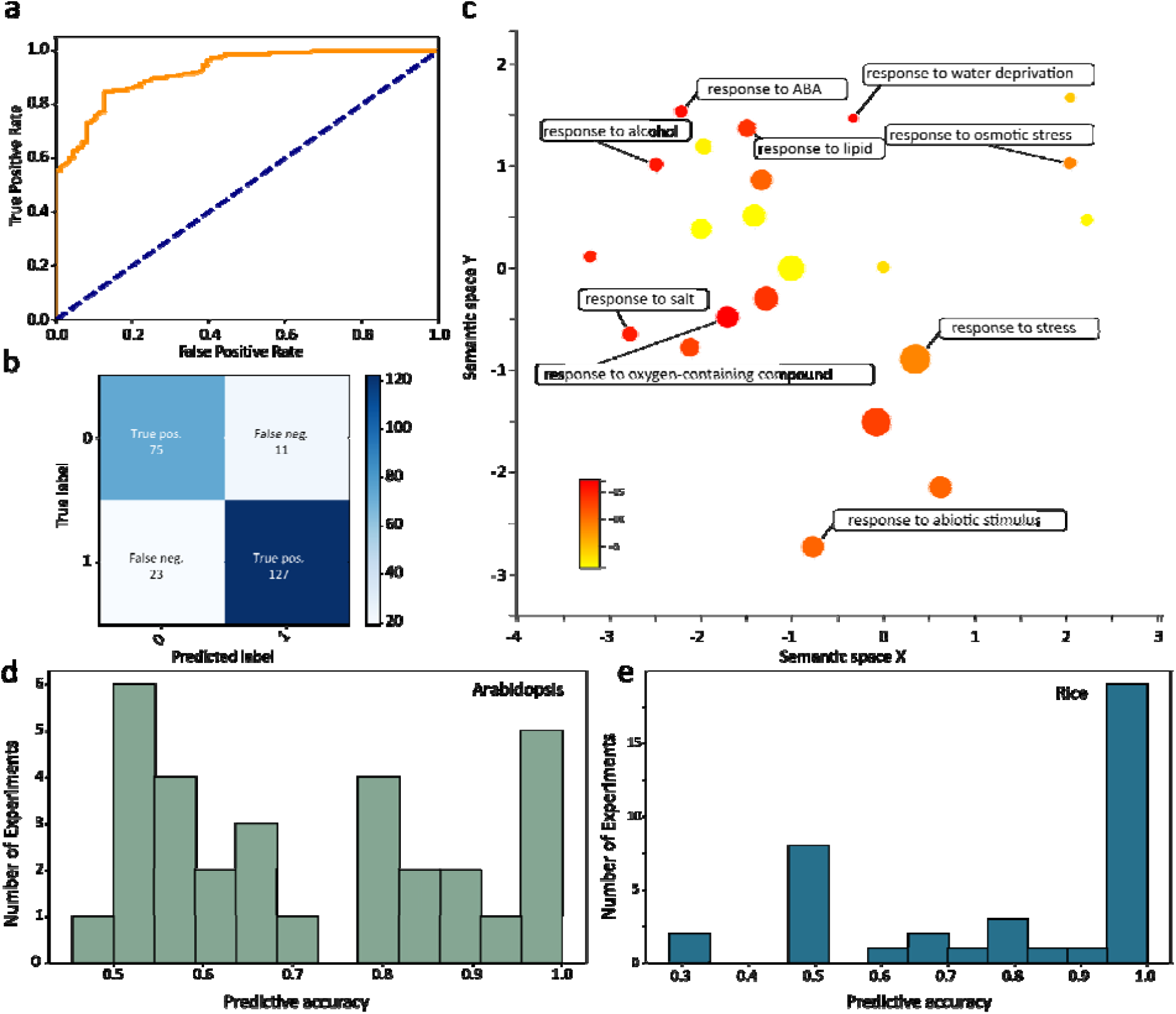
Predictive modeling of water stress in drought expression data. (a) Receiver operating characteristic curve showing the performance of the Random Forest based drought classification model across all classification thresholds. (b) Confusion matrix of the drought predictive model. (c) Multi-dimensional scaling plot showing clusters of enriched gene ontology terms for the top 100 most important features (genes) in the Arabidopsis Random Forest machine learning models. The size of each circle is proportional to the number of genes annotated with each term and the circles are colored by the log10 of the adjusted p-value. Histogram of the predictive accuracy of the Random Forest models for classifying drought using a leave one experiment out approach for Arabidopsis (d) and Rice (e). A predictive accuracy of 1 corresponds to perfect prediction, and 0.5 is more or less random.

To identify the most important underlying features in the model, we developed a second Random Forest classifier using training data from each Arabidopsis BioProject. The overall accuracy improved significantly to 86% with a 0.77 precision and 0.87 recall for control and 0.92 and 0.85 for drought, respectively (Figure 4a, b). Random Forest classifiers rank and quantify the importance of each feature in the underlying testing dataset, and we surveyed which features (genes) were most important for our drought predictive model. The top 100 genes with the most predictive power are enriched in Gene Ontology (GO) terms exclusively related to drought processes (Figure 4c). This includes abscisic acid-activated signaling, and responses to osmotic stress, water stress, salt stress, oxygen-containing compounds, and ABA among others. This is perhaps not surprising, but supports that our model is using genes with well supported roles in drought responses to make its classification. Among the top predictors are regulators of the ABA signaling pathway (HAI1, HAI2, and ABI2), ABA responsive genes (RD29B, DIG2, RAB1), LEA proteins (LEA4-5, ABR, and RAB18), and a lipid transfer protein involved in cuticle formation (LTP3) (Supplemental Table 3). The top predictors also include genes with unknown function (Supplemental Table 3), and predictive modeling may be used to identify new genes with uncharacterized roles in drought stress responses.

## Variability in drought expression profiles across diverse crop and model plants

Our analyses suggested that drought RNAseq data in Arabidopsis is wildly variable, but is this a unique feature of Arabidopsis or a wider issue with drought studies in plants? To answer this question, we re-analyzed published RNAseq data for five additional plant species with between 12-57 individual drought experiments. This includes 179 soybean, 318 tomato, 137 wheat, 1,701 maize, and 981 rice RNAseq samples (Supplemental table 1). Notably, these experiments exhibit greater variability compared to Arabidopsis in terms of genotypic diversity, tissue type, drought assay methods, environmental conditions (e.g., greenhouse, growth chamber, or field settings), and stress severity. We processed this data using the same analytical pipeline that we applied to the Arabidopsis data, utilizing the most recent or highest-quality reference genome available for each species. Similar to the Arabidopsis results, Principal Component Analysis (PCA) did not reveal clear distinctions between drought-exposed and control samples for any of the species (Figure 5). This lack of separation could potentially be attributed to experimental artifacts, and to address this, we applied ComBat to mitigate batch effects across all of the BioProjects. The application of ComBat led to a reduction in variability among the studies for maize and rice, resulting in distinct clustering of drought and control groups in the adjusted expression data (Figure 5). However, when applying ComBat to the wheat, soy, and tomato datasets, we did not observe a clear clustering pattern based on stress level, mirroring our observations in Arabidopsis. We tested Leave-One-Group-Out cross-validation in rice to test if predictive models could more efficiently discern drought and control given the improved sample comparability compared to Arabidopsis. The aggregate performance across all models was 0.82, with a precision of 0.84 and a recall of 0.92. Performance metrics for individual datasets generally exceeded those observed in Arabidopsis, with 18 datasets achieving perfect predictive accuracy (Figure 4E). Together, this suggests datasets outside of Arabidopsis profile drought more consistently and more comparable, but there is still significant variability.

**Figure 5.**
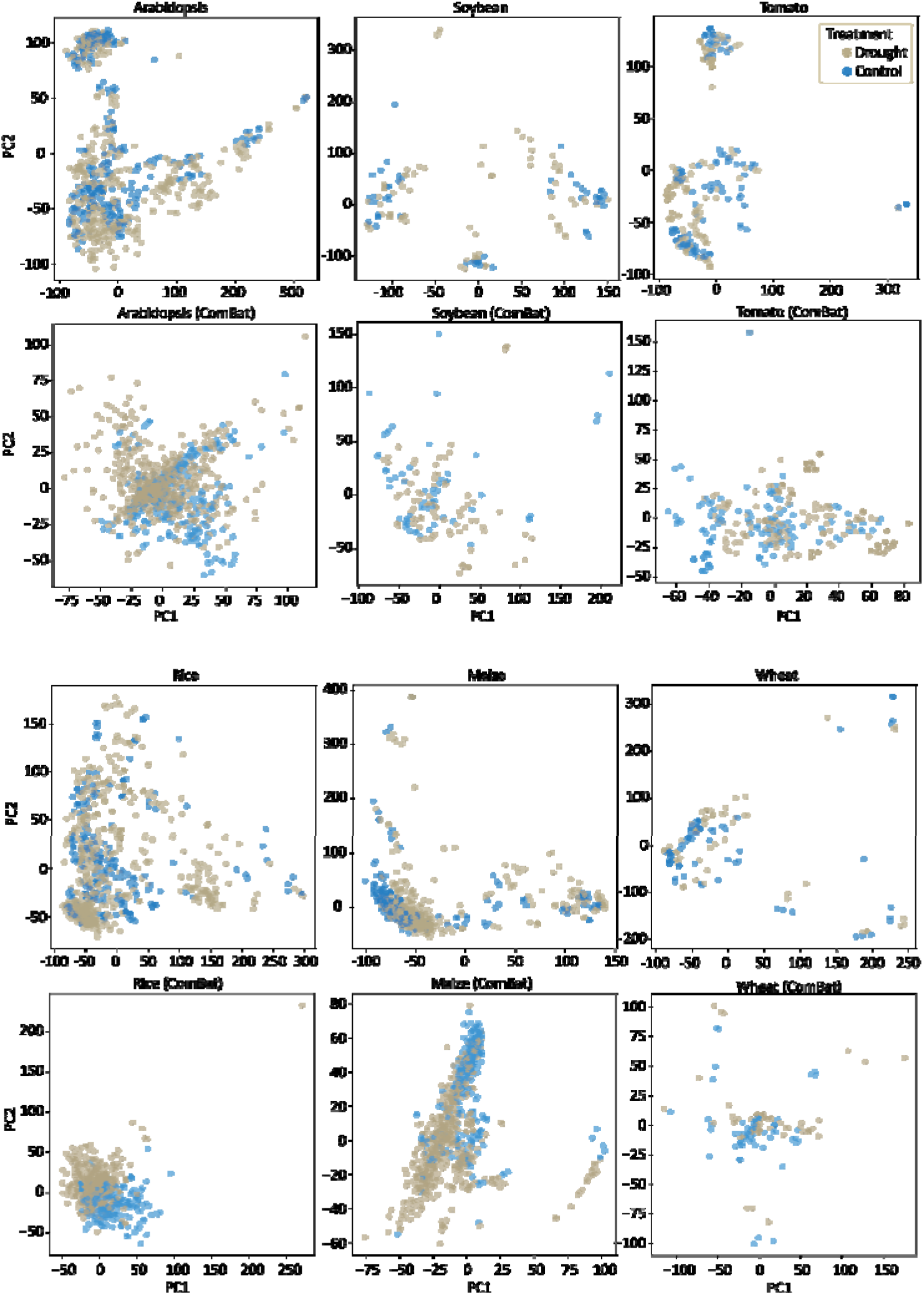
Variability of drought gene expression datasets across model and crop species. PCA plots are shown for drought gene expression datasets in the eudicots Arabidopsis, soybean, and tomato (top) and monocots rice, maize, and wheat (bottom). The PCA of Arabidopsis data from Figure 2a is re-drawn here to enable easier comparison. The first two principal components are plotted for all samples and colored by different factors including a binary classification of drought (blue) and control (gold). Samples corresponding to recovery or unclear timepoints are not included. PCAs from Log2 transformation of the raw TPMs are shown on the top panel for each species, and ComBat corrected samples are shown below.

## Discussion

Genome-scale datasets have increased exponentially, and millions of individual studies from plants are publicly available. These datasets span the genomic, transcriptomic, epigenetic, and chromatin landscapes for thousands of diverse species aimed at numerous research questions related to plant form and function. Historically, most of these datasets were examined in isolation, often due to the specialized nature of each study and the lack of tools or interest in integrative analyses (Sielemann et al., 2020). However, recent computational and methodological advancements encourage data integration into more comprehensive comparative frameworks to identify conserved or emergent features of plant systems (Sreedasyam et al., 2023). A critical aspect of comparing multiple datasets is ensuring uniformity and consistency across experiments. Without this, the nuances of the data can become blurred, as sparsity and heterogeneity have the potential to overshadow crucial biological insights. Here, we re-analyzed the wealth of Arabidopsis drought gene expression data and found that inconsistencies across experiments, potential quality issues, and a lack of paired physiology data complicate integrative analyses. Below, we discuss the factors underlying these issues and propose guidelines that can enable enhanced comparability and reproducibility of future studies.

Applying consistent abiotic stress is relatively straightforward for heat, cold, salinity, UV, light, or nutrient deficiencies. Temperature can be raised or lowered for a set time, plants can be subjected to too much or too little light, and solutes or micronutrients can be maintained to predetermined molarities or concentrations to induce the desired deficiency. These discrete conditions enhance reproducibility across labs and enable cross-study comparisons. Our re-analysis of heat stress data in Arabidopsis supports this claim, as we identified a consistent expression profile under heat regardless of differences in growth conditions, developmental timing, tissue, or other variables across labs and separate studies. Drought stress in contrast, is more difficult to apply, and the lack of standardization makes it difficult to compare experiments or even distinguish between control and stress samples in some cases. Drought itself is a complex and ill-defined stress with unclear delineations of mild and severe or tolerance and avoidance. More than other abiotic stresses, environmental contexts such as the volume and composition of soil or media, temperature, vapor pressure deficit, light intensity, and air flow affect the progression of drought stress. Plants growing in small pots, well-draining soil, higher temperatures, more light, or low humidity will dry faster. Days of drying is reported for most Arabidopsis studies, but this is an arbitrary metric that does not reflect discrete plant physiological states and is impossible to standardize across conditions without additional data to measure the water status of the plant and its soil. Even drought studies performed in the same environment can be challenging to standardize, as different genotypes or mutants can use water at different rates, causing individual plants to experience differing drought severities within the same timeframe and experiment (Ginzburg et al., 2022).

We observed that a surprising 39% of drought RNAseq experiments in Arabidopsis report no drought physiology data, potentially undermining any findings, and preventing these datasets from being used in comparative studies or meta analyses. Many of these studies simply withheld watering for days to weeks and collected control and ‘drought’ stressed tissues for RNAseq and other downstream analyses. No measurements of soil water content or physiological responses were performed. Other studies measured only soil water content without paired measurements in plant tissues, providing no estimate of how stress was affecting the plants. These factors likely explain why we were unable to clearly separate stressed and control samples across studies, even when controlling for experimental artifacts and batch effects. Many ‘drought’ samples have expression signatures that mirror well-watered tissues, suggesting that many plants were not experiencing the physiologically relevant drought stress authors thought had been applied. This noisiness and heterogeneity made it difficult to develop an accurate predictive model of drought stress, and the predictive accuracy was more or less random for almost a third of studies.

Drought datasets outside of Arabidopsis are generally higher quality, with more consistent application of drought conditions and distinct signatures of water deficit responses. The reason for this is unclear. It is not difficult to stress Arabidopsis, and physiological signatures of water stress can be seen with a moderate decrease in water potential (−0.5–1.5 Mp). Compared to Arabidopsis, maize, rice, tomato, wheat, and soybean are generally grown at higher temperatures under higher light intensity, and they have much higher rates of evapotranspiration. When combined with greater plant biomass and the use of small pots, these species may experience drought stress after only a few days without watering in growth chambers and greenhouse settings. In contrast, small or uncrowded Arabidopsis plants may grow well for a week or more between waterings without experiencing water deficit (Ginzburg et al., 2022). Consequently, the community standard of withholding water for 7-10 days for Arabidopsis may be insufficient for stressing plants, and this could explain the prevalence of lower quality datasets of Arabidopsis research.

Applying consistent drought is challenging and efforts of standardizing stress severity have seen mixed success. Automated phenotyping systems can accurately manage soil water content, but they are expensive, have limited scale and flexibility, and plants are still subjected to other environmental fluctuations. While solutes like PEG or sorbitol can help control soil water potential, they may induce unnatural responses in plants (Gonzalez et al., 2022). Rainout shelters or controlled irrigation can help replicate natural drought conditions, but plants are often subjected to other environmental stresses throughout a growing season. Although there is no universally accepted method for conducting drought studies in plants, accurately quantifying water status and associated physiological responses can facilitate the comparison of experiments across different laboratories (Juenger and Verslues, 2022). Developing a deeper understanding of drought responses requires integrating datasets that range from mild to sublethal and from a wide sampling of genotypes, tissues, and conditions. Rather than standardizing drought, we suggest that researchers should collect paired physiology, biochemistry, and morphological datasets at sufficient temporal and spatial resolution to quantify plant health. These traits can first verify that plants are experiencing the desired level of stress prior to expensive sequencing, and serve as features or covariates for integration and re-analysis of multiple datasets.

## Methods

### Assembling a representative catalog of gene expression RNAseq data

We assembled a database of drought RNAseq data in Arabidopsis, soybean, tomato, rice, maize, and rice from the NCBI sequence read archive (SRA). Bulk data was retrieved using a series of drought or heat stress related keywords with the SRA Advanced Search Builder. The following metadata was collected for each experiment: tissue type(s), developmental stage, environment (e.g., greenhouse, field, growth chamber etc.), media type, duration of stress, mechanism of drying, associated physiology datasets, genotype, number of timepoints, and number of replicates. 112 studies had a linked publication in the NCBI metadata and 130 had no associated publication across all 6 species. Similar metadata was retrieved for individual SRA samples along with a binary classification of treatment (drought or control) where possible.

Metadata was retrieved from the SRA and associated publications, but the lack of publications and ambiguity in some labels led to a high degree of missing or sparse metadata for many samples, and our manual annotations were conservative to reduce mislabeling samples for analysis and downstream predictive modeling.

### RNAseq data processing

Raw RNAseq reads were downloaded from the NCBI SRA and quantified using a pipeline to trim, align, and quantify gene expression data (https://github.com/pardojer23/RNAseqV2).

Briefly, sequence adapters were trimmed and a quality check was performed on the raw FASTQ files using the fastp program (v0.23.2) (Chen et al., 2018). The cleaned sequencing reads were then pseudo-aligned to the Arabidopsis TAIR10 (Cheng et al., 2017), maize (*Zea mays* B73 V5) (Hufford et al., 2021), rice (*Oryza sativa Kitaake* v3.1) (Jain et al., 2019), tomato (*Solanum lycopersicum* ITAG4.0) (Hosmani et al., 2019), soybean (*Glycine max var. Williams* 82 V4) (Valliyodan et al., 2019), wheat (*Triticum aestivum cv. Chinese Spring* RefSeq v2.1) (Zhu et al., 2021) genomes using salmon (v1.6) (Patro et al., 2017). The transcript level counts were converted to gene level using the R package TXimport (v 1.22.0) (Soneson et al., 2015). Raw TPMs or log2+1 transformed values were used for downstream analyses. The median alignment rate is 69.1% across all species and 70.8% in Arabidopsis, 79.5% in maize, 62.0% in rice, 65.1% in tomato, 64.7% in soybean, and 64.9% in wheat. These alignment rates are consistent with other meta analyses of gene expression in Arabidopsis (Zhang et al., 2020).

Principal Component Analysis (PCA) was performed using built in functions in Scikit-learn (Pedregosa et al., 2011) on the log2 transformed gene expression data (TPMs) to reduce dimensionality and capture the main sources of variation within the datasets. The first two principal components were plotted for each species and labeled by various factors.

### Predictive modeling of drought stress responses

Our previous work on predictive modeling using drought gene expression in sorghum and maize found that the Random Forest ensemble learning method performed best for classification (Pardo et al., 2022), so we first tested Random Forest on our data. Random Forest models were constructed using the RandomForestClassifier function from scikit-learn (v1.1.0) (Pedregosa et al., 2011). To select the hyper-parameters, the RandomizedGridSearchCV function was utilized, with 100 iterations employing 3-fold cross-validation to traverse the parameter space. Samples were split into 75% training and 25% testing, and model performance was compared using the full unbalanced datasets for each species as well as balanced subsets. Analyses using balanced datasets in Arabidopsis had slightly lower but similar precision and recall, so the full set of samples were used. Feature importance was calculated using the mean decrease in impurity (Gini score) as implemented in scikit-learn (v1.1.0). All genes were subsequently ranked by their respective importance score. Enriched Gene Ontology terms were calculated for the top predictive features in Arabidopsis using the Panther classification system (Mi et al., 2013).

We also tested predictive classification for three additional machine learning algorithms: Linear Support Vector Classifier (LinearSVC), Simple Neural Network (via the Multi-layer Perceptron classifier, MLPClassifier), and Histogram-based Gradient Boosting Classifier (HistGradientBoostingClassifier) implemented using scikit-learn (v1.1.0). The Linear Support Vector Classifier was implemented using the LinearSVC classand the fit method was used to train the model, and the predict method was applied to generate predictions. A simple neural network was developed using the MLPClassifier class. We initialized the MLP with one hidden layer of 100 neurons, and trained the model using the fit method before making predictions with the predict method. The Histogram-based Gradient Boosting Classification Tree, a variant of gradient boosting that is much faster than the traditional Gradient Boosting Classifier, was implemented using the HistGradientBoostingClassifier class from the sklearn.ensemble module. This algorithm is capable of handling missing values, and it also applies the ‘Early-Stopping’ method to avoid overfitting. For all the models, we evaluated their performance by calculating the accuracy score and generating a classification report, which included precision, recall, f1-score, and support for each class. Random Forest outperformed other classifiers in all instances, and was thus used for downstream analyses.

## Supporting information

Supplemental Tables and Figuyres

## Data availability

The data analyzed in this meta-analysis are detailed in Supplemental Table 1, including Sequence Read Archive (SRA) identifiers, PubMed IDs, and all other sample metadata. Raw expression values, expressed in transcripts per million, are accessible on Dryad (https://doi.org/10.5061/dryad.7sqv9s50g). Jupyter notebooks containing all Python code used in this project, along with additional metadata, are available on GitHub: https://github.com/bobvanburen/Drought_meta_analysis_VanBuren_etal_2024.

## Acknowledgements

This work was funded, in part, by the Water and Life Interface Institute (NSF-DBI-2213983) to SYR, RAM, and RV, the United States Department of Agriculture National Institute of Food and Agriculture (USDA-NIFA 2022-67013-36118) to RV, and the US Department of Energy, Office of Science, Office of Biological and Environmental Research, Genomics Sciences Program grants DE-SC0021286 and DE-SC0023160 to SYR. CM, JP, and JS were supported by the predoctoral training award T32-GM110523 from the National Institute of General Medical Sciences of the NIH. AP, JS, and MLW were supported by the National Science Foundation Research Traineeship Program (NSF-NRT 1828149).

## Notes

### Competing Interest Statement

The authors have declared no competing interest.

