## Supplemental Tables and Figuyres for "Variability in drought gene expression datasets highlight the need for community standardization"

**Supplemental information VanBuren et al. 2024.**

**Supplemental Figures**

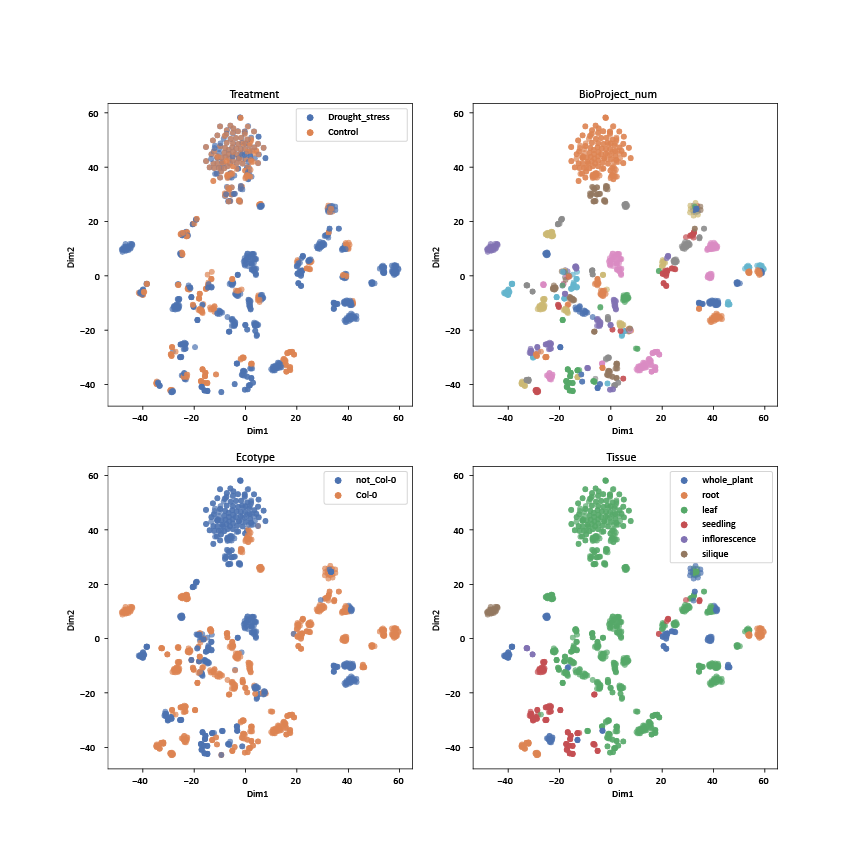

**Supplemental Figure 1. Visualization of Arabidopsis drought gene expression data via t-SNE.** This figure displays the first two dimensions of the t-SNE transformation applied to all samples and colored by different factors including a binary classification of drought and control (upper left), BioProject (upper right), genotype/accession of the sample (Col-0 or others; bottom left), and the tissue type (bottom right).

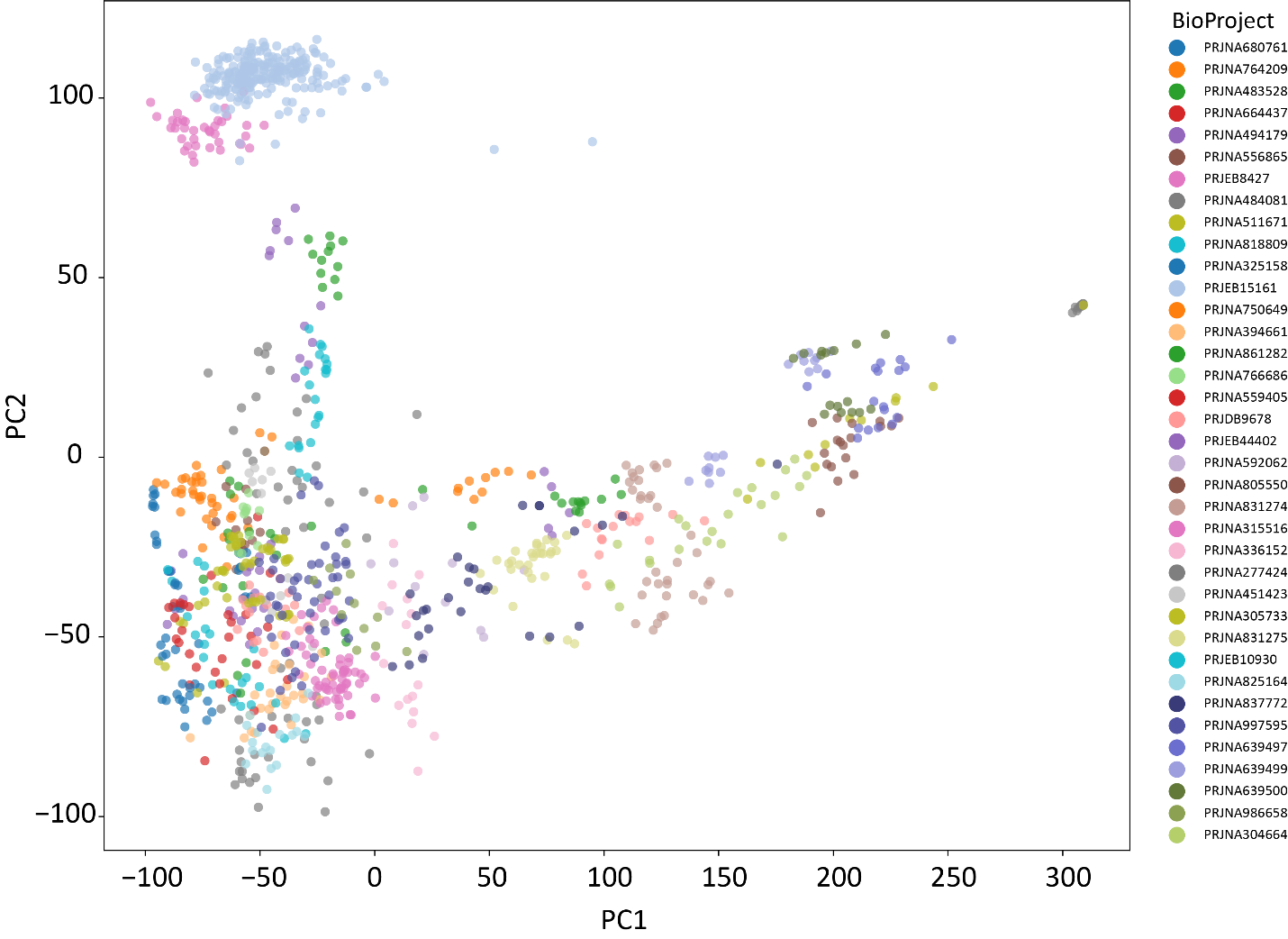

**Supplemental Figure 2. Expanded PCA of Arabidopsis drought RNAseq data colored by BioProject.** The data from Figure 2 is replotted and recolored, but only BioProjects with 12 or more samples are shown in this PCA for simplicity.

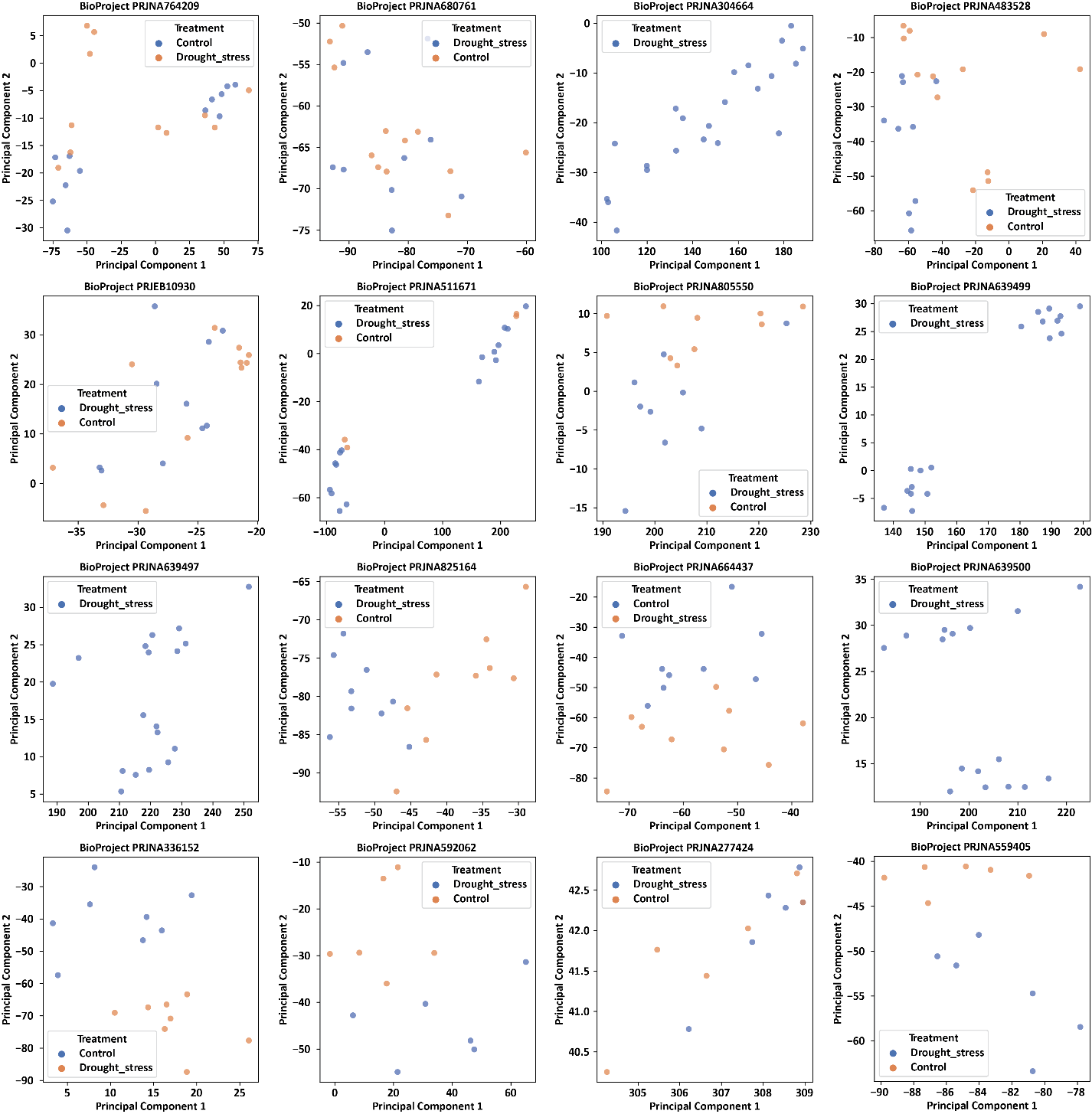

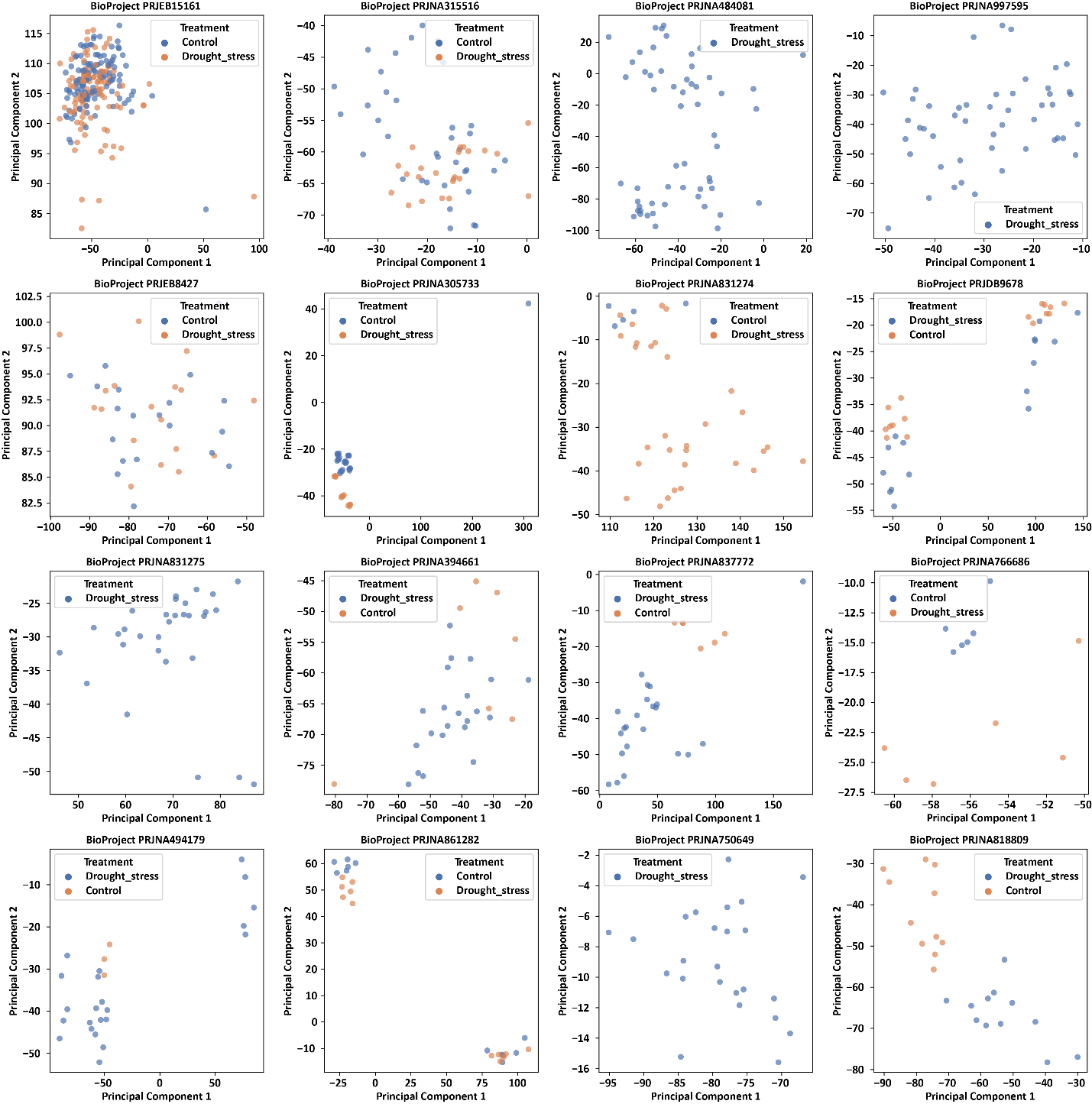

**Supplemental Figure 3. Principle component analysis of Arabidopsis drought data by individual experiment (BioProject).** Only experiments/BioProjects with 12 or more samples are shown.

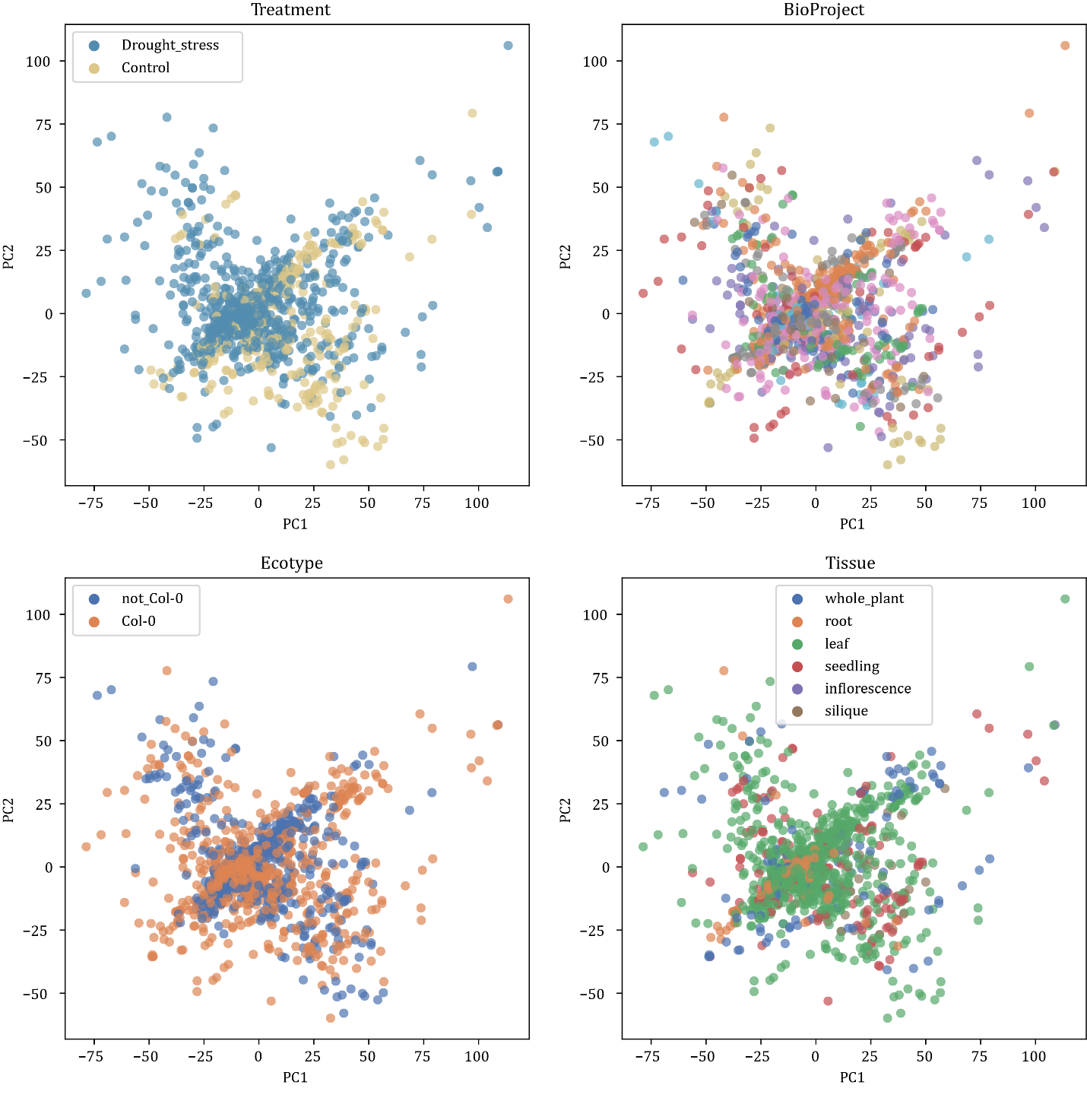

**Supplemental Figure 4. Principal component analysis of Combat adjusted Arabidopsis drought RNAseq data.** The first two principal components are plotted for adjusted expression values and colored by different factors including a binary classification of drought and control (upper left), BioProject (upper right), genotype/accession of the sample (Col-0 or others; bottom left), and the tissue type (bottom right).

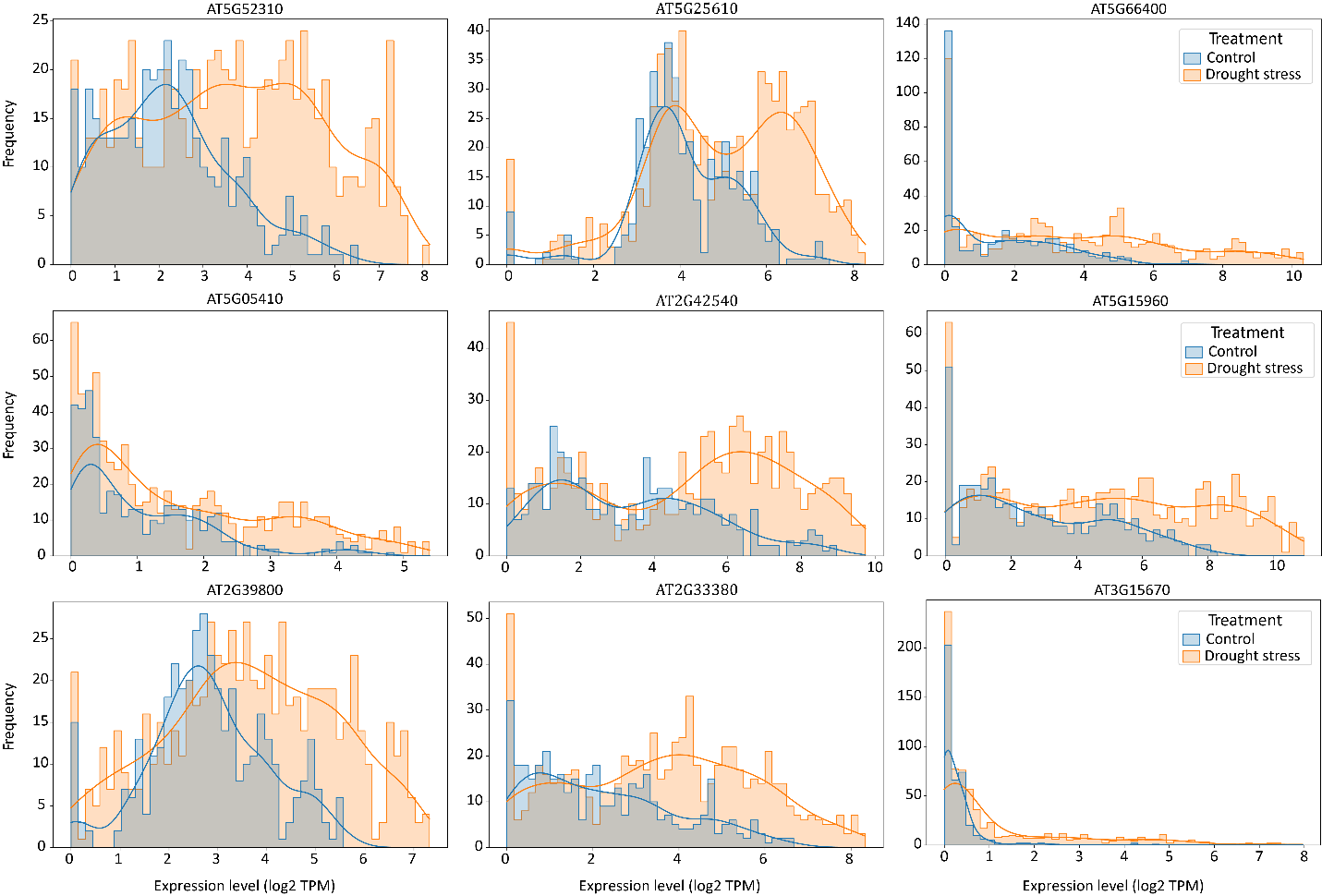

**Supplemental Figure 5. Histogram of drought marker gene expression across Arabidopsis RNAseq data.** Log2 transformed expression values are plotted for each drought (orange) and control (blue) sample for the classic drought marker genes shown in Figure 2.

**Supplemental Tables**

**Supplemental Table 1. Summary of the NCBI BioProjects analyzed in this project. (see external excel file).**

**Supplemental Table 2. Precision and recall values for all classifiers tested using the Arabidopsis drought gene expression data.**

|  | **Random Forest** | **K-Nearest Neighbors** | **Linear Support Vector** | **Multi-Layer Perceptron** | **HistGradientBoosting** |
| --- | --- | --- | --- | --- | --- |
| Precision (control) | 0.6 | 0.49 | 0.49 | 0.52 | 0.59 |
| Recall (control) | 0.84 | 0.88 | 0.93 | 0.96 | 0.83 |
| f1-score (control) | 0.7 | 0.63 | 0.64 | 0.67 | 0.69 |
| Precision (drought) | 0.79 | 0.66 | 0.74 | 0.86 | 0.77 |
| Recall (drought) | 0.51 | 0.2 | 0.16 | 0.22 | 0.49 |
| f1-score (drought) | 0.62 | 0.3 | 0.26 | 0.36 | 0.6 |
| **Overall accuracy** | **0.66** | **0.51** | **0.52** | **0.56** | **0.65** |

**Supplemental Table 3. Top features (genes) from the Random Forest based drought classifier model in Arabidopsis (see separate Excel file).**
